# Utilizing Aggregated Molecular Phenotype (AMP) Scores to Visualize Simultaneous Molecular Changes in Mass Spectrometry Imaging Data

**DOI:** 10.1101/2023.06.01.543306

**Authors:** Jessie R. Chappel, Mary E. King, Jonathon Fleming, Livia S. Eberlin, David M. Reif, Erin S. Baker

## Abstract

Mass spectrometry imaging (MSI) has gained increasing popularity for tissue-based diagnostics due to its ability to identify and visualize molecular characteristics unique to different phenotypes within heterogeneous samples. Data from MSI experiments are often visualized using single ion images and further analyzed using machine learning and multivariate statistics to identify *m/*z features of interest and create predictive models for phenotypic classification. However, often only a single molecule or *m/z* feature is visualized per ion image, and mainly categorical classifications are provided from the predictive models. As an alternative approach, we developed an aggregated molecular phenotype (AMP) scoring system. AMP scores are generated using an ensemble machine learning approach to first select features differentiating phenotypes, weight the features using logistic regression, and combine the weights and feature abundances. AMP scores are then scaled between 0 and 1, with lower values generally corresponding to class 1 phenotypes (typically control) and higher scores relating to class 2 phenotypes. AMP scores therefore allow the evaluation of multiple features simultaneously and showcase the degree to which these features correlate with various phenotypes, leading to high diagnostic accuracy and interpretability of predictive models. Here, AMP score performance was evaluated using metabolomic data collected from desorption electrospray ionization (DESI) MSI. Initial comparisons of cancerous human tissues to normal or benign counterparts illustrated that AMP scores distinguished phenotypes with high accuracy, sensitivity, and specificity. Furthermore, when combined with spatial coordinates, AMP scores allow visualization of tissue sections in one map with distinguished phenotypic borders, highlighting their diagnostic utility.

## INTRODUCTION

The accurate identification of different tissue phenotypes is crucial for early diagnosis, successful tumor removal, and treatment of disease. Achieving this has been challenging due to the spatially complex nature of tissues, especially tumors, which gives rise to both intra-tumoral and inter-tumoral heterogeneity.^1, 2^ Historically, the manual evaluation of stained or labeled sections of tissue by highly trained histopathologists has been the gold standard for diagnosis. However, hematoxylin and eosin (H&E) stains only provide morphological information and may not fully resolve tissue types, especially when changes are primarily molecular in nature.^3, 4^ Furthermore, this approach can also be time-consuming, subjective, and ultimately delay patient care. Therefore, the need for an improved approach that combines molecular and spatial distributions with tissue morphology has become apparent. Mass spectrometry imaging (MSI) has thus emerged as a powerful approach to address these needs.^5–9^

MSI-based techniques rely on sampling regions of interest by using an ionization probe that feeds directly to a mass spectrometer. Notably, ambient ionization sampling techniques allow for the direct analysis of complex samples under atmospheric pressure and require minimal sample preparation, thereby allowing for rapid data collection.^6, 10^ To identify intrinsic patterns in the resulting datasets, unsupervised statistical approaches are often applied, which utilize the *m/z* features detected to assess trends without prior biological knowledge. Due to the large size of MSI datasets, it is common to implement dimension reduction techniques, such as principal components analysis (PCA), t-distributed stochastic neighbor embedding (t-SNE), or non-negative matrix factorization (NMF) as a pre-processing step. Dimension reduction is often followed by clustering techniques, such as hierarchical clustering, k-means clustering, or Gaussian mixture modeling to segment pixels into groups with similar feature profiles. Once clusters are assigned, tissue sections are often visualized by creating ion images where pixels are colored according to their cluster. Comparison of cluster localization to pathologist annotated slides has revealed that these approaches can identify highly relevant clusters that correspond to regions of the tissue that have different biochemical or biological properties, such as different cell types or different stages of disease.^11–18^

While unsupervised approaches aim to unveil natural patterns in the data, supervised approaches are often used to select features associated with phenotypes of interest. This can be achieved by either applying univariate methods, such as an ANOVA or t-test, to directly characterize the relationship between a given feature and phenotype, or by using multivariate approaches, such as random forest (RF) or support vector machine (SVM), to highlight features that are important for phenotypic classification.^19,20, 21^ For example, supervised approaches have been used to identify diagnostic lipid and metabolic signatures of human cancerous tissues including brain^22^, breast^23, 24^, thyroid^25^, gastric^26^, ovarian^27^, and others^19, 28-30^. With these approaches, ion images are typically made by coloring pixels based on feature abundance for features that were found to be statistically significant or influential on model performance. Alternatively, tissue sections can be visualized with a single image by coloring pixels based on phenotypic predictions provided by models.^26, 31^

Although the benefits of using MSI data for tissue diagnosis and visualization are apparent, there are still limitations in the outlined statistical approaches. While unsupervised techniques have the benefit of requiring minimal *a priori* knowledge, resulting clusters may not correspond to phenotypic differences. Further, because pixels are colored categorically, rigid borders will exist between clusters in ion images, failing to capture any gradual molecular changes. These limitations may be circumvented by utilizing a supervised approach, where specific differences between phenotypes can be assessed based on sample annotations and tissue sections are visualized based on feature abundances. However, because often only one feature is plotted at a time, this approach results in a large number of images to examine, which is not only time consuming, but also makes it difficult to holistically understand the molecular landscape of a tissue section. Moreover, while supervised analyses may produce accurate predictive models for some phenotypes, they typically only provide categorical predictions, eg, normal or cancer, which lack nuance. For example, while a histopathologist may identify unusual or rare features in a tissue section, a predictive model may only output that the tissue is normal with no further indication of anomalies that may be relevant to disease management and prognosis, eg, detection of inflammation or precancerous lesions.^32^ To this end, we have developed aggregated molecular phenotype (AMP) scores to allow concurrent visualization of phenotype associated features and improve the accuracy and nuance of predictive models by having a continuous rather than categorical outcome.

An AMP score is a value between 0 and 1 that summarizes the aggregated molecular information of a single sample or pixel in an MSI experiment. These scores are defined for pairwise phenotype comparisons with low scores indicating a given pixel has molecular characteristics matching the user-defined class 1 phenotype (often control) and high scores correlating with the user-defined class 2 phenotype. To calculate AMP scores, we utilized an ensemble feature selection approach to identify features differing between the two phenotypes, weighted these features using logistic regression, and then applied a custom function to combine feature abundances with their respective weights for each sample or pixel. A threshold value of 0.5 was utilized for binary classification of these scores with scores closer to 0.5 suggesting that a pixel shares molecular characteristics of both phenotypes. Once calculated, AMP scores were then combined with the location coordinates for each pixel to create one image of a tissue section. This image represents all features (or molecules) of interest and illustrates how the molecular landscape changes spatially (Figure 1).

**Figure 1:**
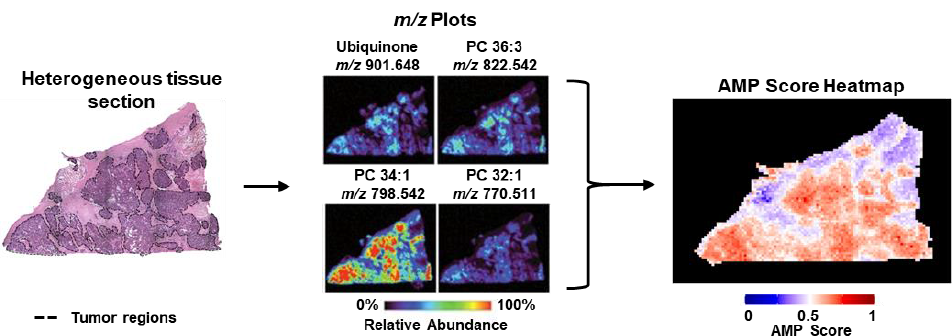
Example AMP score heatmap for a heterogeneous tissue sample. The left shows an H&E stain for a high-grade serous carcinoma tissue sample with tumor areas outlined in dashed black lines. With current MSI approaches, feature abundances across the tissue area are visualized individually, resulting in many plots (middle).^27^ However, using AMP scores (right), the same tissue section is visualized by summing across all features weights and abundances to greatly reduce the number of images needed.

In this paper, we utilized metabolomic data from three previously published studies to demonstrate how AMP scores are calculated and evaluate their performance.^23, 25, 27^ Namely, in the first evaluation, we analyzed AMP scores for homogenous tissue sections from follicular thyroid adenoma (FTA), a benign thyroid tumor, vs. papillary thyroid carcinoma (PTC),^25^ the most common type of thyroid cancer. In our next comparison, we assessed normal breast (NB) vs. invasive ductal carcinoma (IDC), the most common type of breast cancer tissue to evaluate margins.^23^ Three different ovarian tissues, normal ovarian tissue (NO), borderline ovarian tumor (BOT), and high-grade serous carcinoma (HGSC) were evaluated in the final comparison to understand how each pairwise assessment would perform, especially when our class 1 was not a control but a less severe disease case (ie, BOT vs HGSC)^27^. These analyses illustrated the high predictive power of AMP scores with class 1 samples having substantially lower scores compared to the class 2 samples. The ability to distinguish phenotypes was further showcased in AMP score heatmap visualizations, which distinguished tissue borders and identified regions of normal and diseased tissue. Overall, these results suggested that AMP scores can be used as a powerful approach to visualize and diagnose tissue sections from MSI.

## METHODS

### MSI Datasets

AMP scores were constructed and validated using metabolomic MSI data from three previously published studies.^23, 25, 27^ These studies evaluated frozen human tissue sections using desorption electrospray ionization (DESI) MSI to characterize the molecular profile of the different phenotypes. In this work, we considered a subset of features from negative ion mode data from each study, with each feature having an *m/z* value, associated abundance, and x and y coordinates, associated with the MS image. In total, 210 tissue samples were analyzed including: 20 follicular thyroid adenoma (FTA) (15,088 pixels), 20 papillary thyroid carcinoma (PTC) (13,554 pixels), 41 normal breast (NB) (1,674 pixels), 81 invasive ductal carcinoma breast cancer (IDC) (31,639 pixels), 1 heterogeneous NB/IDC (1,910 pixels), 15 normal ovarian (NO) (11,126 pixels), 14 borderline ovarian tumor (BOT) (3,759 pixels), 14 high grade serous carcinoma (HGSC) (9,965 pixels), and 2 heterogeneous NO/HGSC (7,304 pixels) samples. Five pairwise phenotype comparisons were then assessed in the study: (1) FTA vs. PTC samples from DeHoog *et al.*, (2) NB vs. IDC samples from Porcari *et al*., and (3) NO vs. BOT, (4) NO vs. HGSC, and (5) BOT vs. HGSC from Sans *et al*. Further information on data collection can be found in their respective publications.^23, 25, 27^

### Overview of AMP Score Calculation Pipeline

AMP scores were calculated by first filtering and normalizing the data. The data were then split into training and testing sets, where distinguishing features were selected and AMP score parameters were calculated using the training data. Finally, AMP scores were calculated for the testing data for evaluation. An overview of this pipeline is shown in Figure 2 and each individual step is described below.

**Figure 2:**
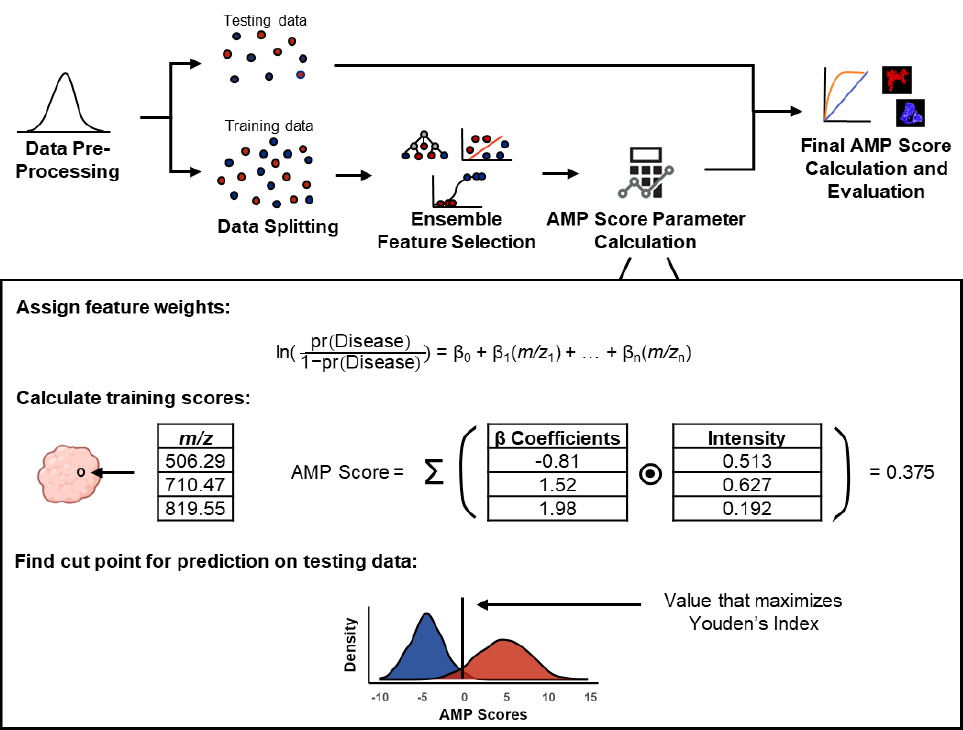
Pipeline for AMP score calculation. Data are first filtered and normalized before being split into testing and training sets. Features distinguishing phenotypes are selected using ensemble feature selection with Lasso regression, random forest, and support vector machine. Selected features are weighted using logistic regression and preliminary AMP scores are calculated for each pixel in the training data by multiplying the abundance of selected features by their respective weights and then summing. Following preliminary scoring, the value that maximizes Youden’s index is calculated, which is later used to scale the testing data. Finally, AMP scores are calculated for the testing data, predictions are made, and scores are plotted for visualization across tissue sections.

### Data Pre-Processing

For each pairwise comparison, data were filtered by first binning each *m/z* value to the nearest hundredth and removing *m/z* values that appear in fewer than 10% of pixels. For all thyroid and ovarian tissue samples, the *m/z* range was restricted to 100-1200. Breast tissue data was collected across multiple laboratories, and cosine similarity analyses of the full mass spectra obtained from the different labs revealed that a higher m/z range was necessary to achieve high spectral similarity. The *m/z* value range for the breast tissue samples was thus restricted to 700-1200 as done in the original manuscript to maintain consistency.^23^ Once all unique features were identified, missing values were imputed using the minimum detected value in the dataset. Following filtering, all abundances were log_2_ transformed and normalized at the sample level by dividing by the peak with the highest abundance value.

### Data Splitting

To evaluate the performance of the AMP scores, we followed standard practice of data splitting in machine learning by first training the scoring system on a subset of the data and then applying it to an independent testing set. For each pairwise comparison, homogeneous samples were randomly split into training and testing sets using a 2:1 ratio. Heterogeneous samples were withheld as an additional validation set.

### Ensemble Feature Selection

To identify features that distinguish between phenotypes, we leveraged an ensemble feature selection approach. In this approach, features were initially selected by applying least absolute shrinkage and selection operator (Lasso) regression, random forest (RF), and support vector machine (SVM) to the training data. Details of these methods, associated parameters, and their implementation are below:

1. Lasso regression is a method that selects features by imposing a penalty on the size of the regression coefficients, shrinking them towards zero, and resulting in a sparse model that retains only the most important features.^33^ Lasso analysis was performed in R (v4.2.1) using the glmnet package.^34^ For each pairwise comparison, the optimal value for lambda, which controls the strength of the regularization applied, was obtained using 5-fold cross-validation with the function ‘cv.glmnet’. After selecting the lambda value, Lasso logistic regression was performed using the ‘glmnet’ function and all features with a non-zero coefficient were retained.
2. RF ranks features based on how much their inclusion in a forest of decision trees decreases the impurity of predictions quantified using the Gini Index, with features yielding the highest decrease in impurity being considered the most important.^35^ To determine these features, RF models were constructed for each pairwise comparison using the randomForest package in R. Each model was built using the function ‘randomForest’ and consisted of 1,000 decision trees where the number of variables to use as candidates at each split point was equal to the square root of the number of features.^36^ Once a model was constructed, the out of bag error rates were noted and features were ranked using the ‘importance’ function. Low ranking features were then removed from the data and the model was reconstructed. This process was repeated iteratively until overall model performance began to decrease, suggesting all current features were useful for distinguishing phenotypes.
3. SVM with a linear kernel was employed for feature selection by examining the coefficients of the decision boundary, indicating the importance of each feature in predicting the classification outcome.^37^ SVM analysis was conducted using scikit-learn in Python (v3.9.7).^38^ Models were first constructed using the ‘SVC’ function with a linear kernel and C = 1. To evaluate model performance, 5-fold cross-validation was performed on the training data using the ‘cross_val_scores’ function and top features were identified using the ‘coef_’ function. Low ranking features were then removed from the data and the model was reconstructed. This process was repeated iteratively until model performance began to decrease, suggesting all current features were useful for distinguishing phenotypes.

Features were determined to be important and included in AMP score calculation if selected by at least two of the three models.

### AMP Score Parameter Calculation

After selecting significant features that distinguish phenotypes of interest, the next step was to assign a weight to each feature. To achieve this, training data was filtered down to just the selected significant features and logistic regression was performed in R using the ‘glm’ function with the abundances of features as independent variables and the phenotype group as the dependent variable. From this model, each feature was assigned a β coefficient. Preliminary AMP scores were then calculated for each individual pixel by multiplying the β coefficient of each selected feature by the feature’s abundance and then summing all of these values. From these preliminary AMP scores, the optimal threshold value for distinguishing the two phenotypes was chosen by finding the value that maximized Youden’s Index, which is defined as the sum of sensitivity and specificity minus 1.^39^ This was done using the ’cutpointr’ function from the R package cutpointr, which utilizes bootstrapping methodology to identify the value that maximizes a given metric.^40^

### Final AMP Score Calculation and Evaluation

Unscaled AMP scores for testing data were calculated for each pixel using the same approach as the training data, which included multiplying the β coefficient of selected feature per pixel by their associated abundance and then summing all significant features. Testing scores were then scaled between 0 and 1, with all scores below the optimal threshold value determined by the training data to range between 0 and 0.5 and all scores above the threshold value to occur between 0.5 and 1. All pixels with a score less than 0.5 were predicted to be from class 1 samples, while pixels with a score greater than or equal to 0.5 were predicted to be from class 2 samples. Predicted phenotype labels were then compared to true pixel labels from pathology, and accuracy, sensitivity, and specificity were calculated using functions from the R package caret. Receiver operator characteristic curves were also calculated using the ‘roc’ function from the R package pROC.^41^ Violin plots, boxplots, and AMP score heatmaps were then made in R using the package ggplot2 to compare results.^42^ Additionally, AMP score heatmaps were created by first plotting each pixel according to its x and y coordinate and then coloring based on AMP score value.

## RESULTS & DISCUSSION

AMP scores were developed in this work to enhance MSI data visualization by assessing multiple features and molecular changes simultaneously (Figure 1), including gradation of borderline phenotypes, instead of forcing categorical assessments. Assigning AMP scores for datasets of interest had several main steps as shown in Figure 2. These steps included: (1) removing noise and normalizing the data, (2) splitting the data into training and testing sets, (3) applying ensemble feature selection to select significant features (or molecules) distinguishing the phenotypes of interest from the training data, (4) using selected features to calculate AMP score parameters, and (5) leveraging parameters from the training data to calculate AMP scores for the testing data and applying the phenotypic predictions to individual pixels. These steps are all detailed in the Methods section.

As AMP scores are calculated solely on selected features, it is imperative to implement a robust feature selection method. For this reason, we opted for an ensemble feature selection approach, which was preferred due to its ability to overcome the limitations and biases associated with individual selection methods.^43^ In our ensemble we included Lasso, RF, and SVM, as they each have different underlying assumptions and therefore are likely to capture unique patterns in the data.^33, 35, 37, 43^ A breakdown of overlap between the three methods is shown in **Figure S1**. While these selection approaches are able to identify distinguishing features for this study, other selection methods, such as elastic net or ridge regression, may be better suited for different comparisons, and could also be easily incorporated into the AMP score pipeline. Moreover, incorporating additional selection methods into the feature selection ensemble may result in a more stable feature list by ameliorating the biases of individual selection methods.

Once features were selected, weights for the selected features were determined. Because a key component of AMP score calculation is multiplying feature abundances with their respective weights, it was crucial that feature weights were directional, so that high feature abundances associated with the class 1 or control phenotype result in lower AMP scores, while high abundance features correlating with the class 2 phenotype provide higher AMP scores. To achieve this, we assigned β coefficients to each feature using logistic regression. This method was preferred over other feature weighting methods, such as information gain, due to the weights being signed and easily interpretable.^44^ However, we do recognize that some datasets may not meet the underlying assumptions for logistic regression, and consequently other weighting methods may be more appropriate. As such, identifying a nonparametric way to generate signed weights is a focus of future work.

After β coefficients were assigned to each feature, we subsequently calculated preliminary AMP scores for the training data. The purpose of doing this was to identify the optimal cutoff score for distinguishing the two phenotypes in each comparison, which would later be used to scale the AMP scores for the testing data. To choose a cutoff, the score value that maximized Youden’s Index was chosen. This specific statistic was maximized over other potential performance measures, such as accuracy, because it considers both the true positive rate (sensitivity) and the true negative rate (specificity), making it a comprehensive measure of dichotomous diagnostic performance.^45, 46^ Once all parameters were determined, AMP scores were calculated for the testing data. To do this, the testing data was first filtered down to the selected features. The abundance for each feature was multiplied by its respective weight at each pixel, and all resulting products were summed. However, since AMP scores ranges varied for the different comparisons, we scaled all AMP scores between 0 and 1, with the cutoff value determined from the training data set to 0.5. This allowed for easier, consistent interpretation both within and across comparisons.

To assess the AMP score calculations, three different MSI studies with normal, benign, and cancerous human tissue sections were evaluated (Figure 3). Specifically, the first study compared patients with follicular thyroid adenoma (FTA) to those with papillary thyroid carcinoma (PTC) to understand how AMP scores performed when comparing homogeneous tissue regions to one another. Next, normal breast (NB) and invasive ductal carcinoma (IDC) samples were evaluated to determine AMP score performance with heterogeneous tissue sections when both normal and tumor regions were present in the same sample. Finally, a study with three different ovarian tissue phenotypes was evaluated (normal ovarian (NO), borderline ovarian tumor (BOT), and high-grade serous carcinoma (HGSC)). From this study, we compared NO to BOT and NO to HGSC to assess the ability of AMP scores to distinguish normal vs. diseased tissues regions and identify phenotypic margins. We also compared BOT to HGSC to assess how well AMP scores could differentiate between different disease severities or classes. A breakdown of the samples used to construct the AMP score models and their performance is given in Figure 3.

**Figure 3:**
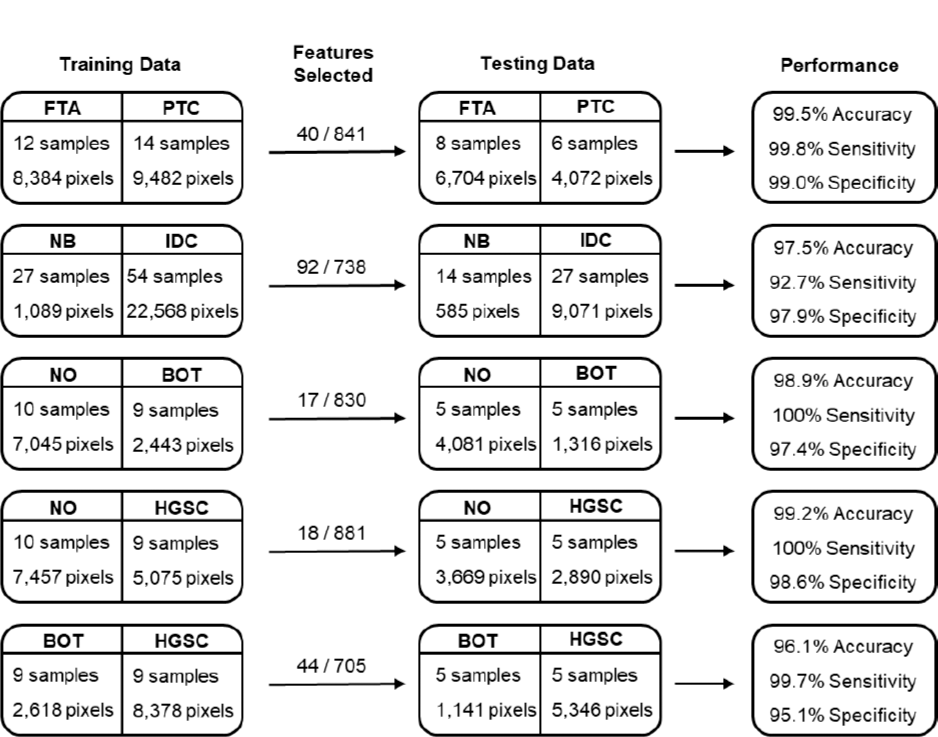
Summary of data used for the AMP score calculation and validation, and overall performance for each pairwise comparison. The number of features selected as significant for each study is also noted versus the total number of features evaluated. From top to bottom: Follicular thyroid adenoma (FTA) vs. papillary thyroid carcinoma (PTC), normal breast (NB) vs. invasive ductal carcinoma (IDC), normal ovarian (NO) vs. borderline ovarian tumor (BOT), NO vs. high grade serous carcinoma (HGSC), and BOT vs. HGSC.

For all studies, the number of pixels used for training far exceeded the number of input features (*e.g.,* 23,657 pixels and 738 features in the NB vs. IDC comparison). This data characteristic supported the decision to perform feature selection using machine learning methods (Lasso regression, RF, and SVM). It was also observed that following ensemble feature selection, the dimensions of each dataset decreased substantially, with 87.5% to 98.0% of the overall features removed prior to AMP score calculation. This reduction resulted in 40 features for the FTA vs. PTC tissue comparison, 92 for NB vs. IDC, 17 for NO vs. BOT, 18 for NO vs. HGSC and 44 for BOT vs. HGSC. When the AMP scores were calculated from these features and assessed on the testing data, they showcased a high predictive power, with class 1 samples having significantly lower scores than class 2 samples. We also observed high sensitivity and specificity across all studies with all metrics above 92.7%, which is visualized in receiver operating characteristic curves (Figure 3 **and Figure S2**). This balance of sensitivity and specificity was observed across comparisons having markedly different pixel ratios. For example, the NB vs. IDC comparison involved considerably more IDC pixels than NB. This is a result of many NB tissue samples being primarily composed of fat, limiting the number of pixels that could be extracted from epithelial cells.^23^ Despite this imbalance, AMP performed comparably to other comparisons. These results suggest that a generalized AMP pipeline offers balanced performance. Importantly, the parameters could be tuned to favor sensitivity (i.e. disease detection) or specificity, depending upon clinical considerations regarding consequences of false-negatives versus false-positives.

While the ability to differentiate phenotypes can be summarized as approximately equal across comparisons as shown by the metrics in Figure 3, the distribution of AMP scores across pixels varied which can be expected based on changes in the tissues studied (Figure 4). In each comparison, the distributions were separated by phenotype, with the greatest separations in AMP scores between the NO vs. HGSC and NO vs. BOT comparisons. Interestingly, in the normal/benign vs. cancerous comparisons (top four plots), the distribution for cancerous pixels was wider than the distribution of pixels for the normal/control This wider distribution may be due to diseased samples having different degrees of progression and thereby leading to more variable molecular profiles compared to control samples. In particular, the distribution of BOT samples appears somewhat bimodal. Since BOTs have the potential to develop into low-grade serous carcinoma, we hypothesize this split may correspond to samples that are progressing in severity.^27^ This trend is further supported in the BOT vs. HGSC distributions (bottom plot), where we see BOT is once again bimodal, with the smaller mode having AMP scores closer to HGSC. When comparing the AMP score distribution of HGSC in the NO vs. HGSC comparison to the BOT vs. HGSC comparison, we also can see that the distribution is much narrower in the BOT vs. HGSC comparison. This result may suggest that the molecular features that differentiate HGSC samples from BOT samples are expressed more consistently in HGSC samples than the features that differentiate NO and HGSC samples. While the comparisons in this manuscript mainly focused on various healthy and disease phenotypes from human tissue, the BOT vs. HGSC comparison showed that the AMP score pipeline is not specific to normal/benign vs. disease cases. Therefore, we believe that AMP scores may be useful for assessing disease severity, identifying mechanisms of pathogenesis, and selecting prognostic markers. To further explore this capability, future steps include characterizing the relationship between patient metadata and AMP scores to determine if correlations exist between scores and patient outcomes or other clinical measurements.

**Figure 4:**
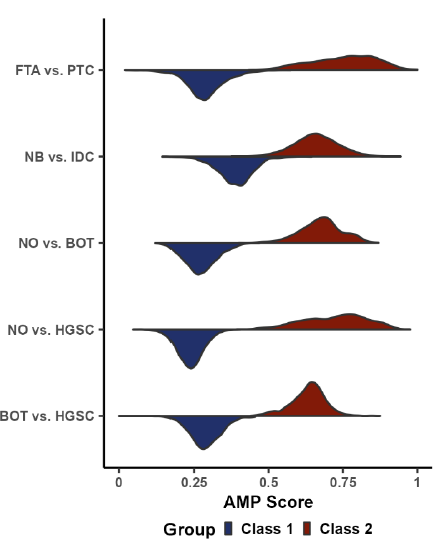
Violin plot showing the density of AMP scores for each pairwise comparison split by phenotype.

To gain more information on the phenotype classifications, we also evaluated the AMP score distributions at the individual sample level (Figure 5). Again, there was a strong distinction between the different phenotypes with samples from the class 1 phenotypes being closer to zero and class 2 samples closer to one. Interestingly, all misclassified pixels had an intermediate AMP score, with the lowest score from the true class 2 pixels being 0.374 and the highest score from the true class 1 pixels being 0.645 (Figure 5A-5E). Because many of the normal/benign tissue samples were extracted from disease adjacent regions, and vice-versa, it is possible that some of these outlier points may represent pixels in the transition region between tissue types and consequently have molecular features like both tissues. An exception to this pattern is sample 7 in the NO/HGSC comparison (Figure 5C), which had over half of the pixels misclassified. This sample only had 20 pixels and therefore very little influence on the overall prediction performance of the testing data. However, this result may also suggest that this sample should be further examined. Additionally, when comparing the sample distributions within each pairwise phenotype comparison to one another, there is considerable variation, particularly for the class 2 phenotypes. This reveals that rather than reflecting the overall phenotype distributions from Figure 4, individual samples or pixels may contribute more heavily to certain parts of the distributions. This trend is best observed when considering the HGSC data in the NO vs. HGSC comparison, where the distribution of all pixels is quite wide (Figure 4), but the individual sample distributions are much narrower and almost completely in distinct interquartile ranges (Figure 5C).

**Figure 5:**
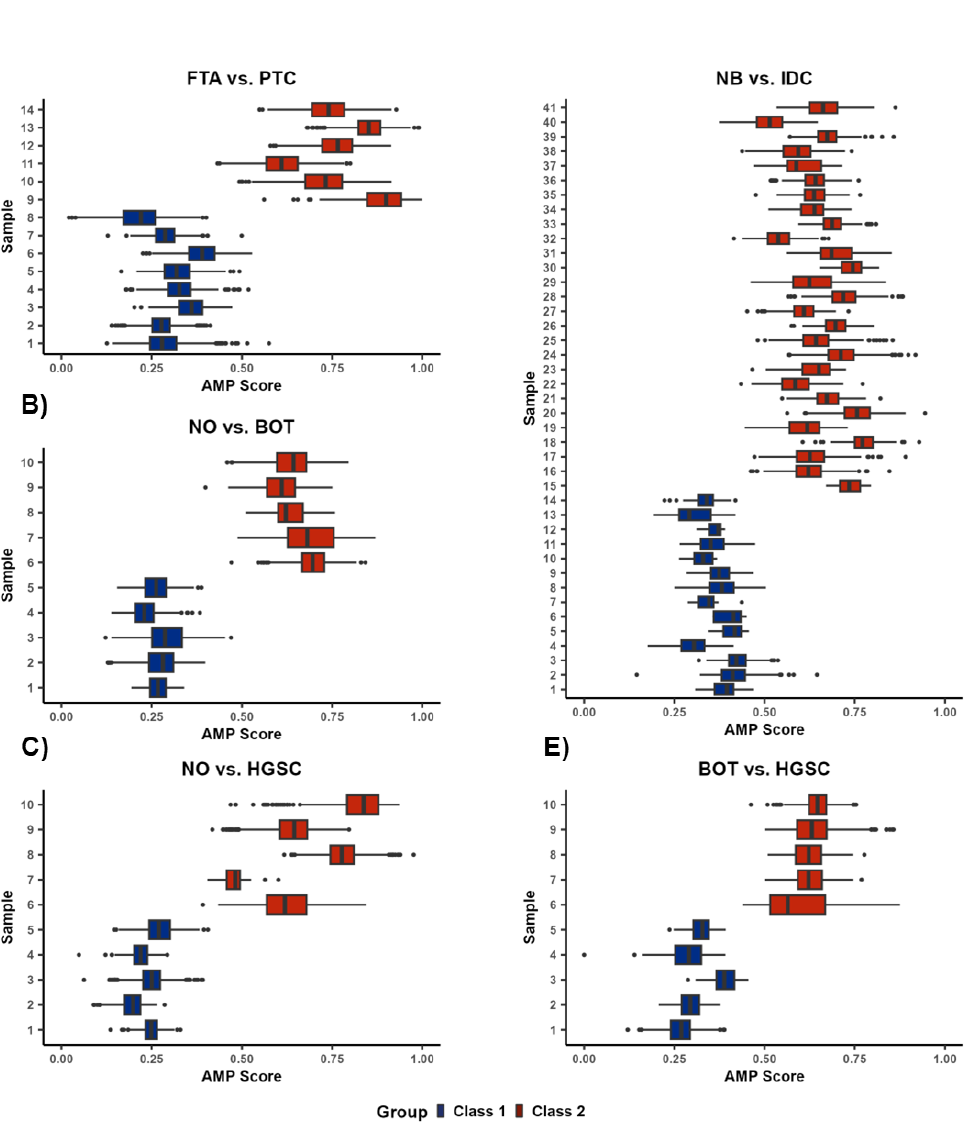
AMP scores distributions by sample for A) FTA vs. PTC, B) NO vs. BOT, C) NO vs. HGSC, D) NB vs. IDC, and E) BOT vs. HGSC. Each individual box and whiskers plot shows the distribution of AMP scores across all pixels within a sample. Outlier points are defined as observations more than 1.5 times the interquartile range away from the upper or lower quartile.

Beyond binary classifications, AMP scores were also evaluated for their continuous scoring of phenotypes and visualization of margins in the tissue sections. The ability of AMP score heatmaps to capture overall molecular patterns was demonstrated in both the homogenous (Figure 6) and heterogeneous samples (Figure 7). In homogeneous tissue sections, the phenotype of each sample was readily apparent from the AMP score heatmaps (Figure 6). For both class 1 and class 2 samples for the FTA vs. PTC, NO vs. BOT, and BOT vs. HGSC comparisons, AMP scores remain relatively consistent throughout the entire diseased or control tissue areas. Notably, pixels with intermediate AMP scores are frequently located near the tissue border, providing further support that outlier points seen in the boxplots may correspond to pixels near phenotypes borders, potentially explaining molecular differences. To further illustrate the ability of AMP scores to differentiate phenotypes, predictions were made on a few heterogenous tissue samples (Figure 7). Specifically, we assessed two HGSC tissues and one IDC tissue, each having accompanying H&E slides with tumorous areas outlined in black. Comparing AMP score heatmaps to these annotations, we see that AMP score heatmaps were able to successfully identify tumorous areas, highlighting the utility of AMP scores for distinguishing phenotypes. Once again, in all three plots, pixels located at the border of normal and cancerous had intermediate scores, suggesting that molecular differences between phenotypes occur gradually, with transition regions sharing molecular characteristics of both tissue types (Figure 7D). This result is consistent with the work done by Woolman *et al.*, which showed that molecular borders are not as sharp as the morphometric borders identified with microscopy, and that a gradient of cancer-like metabolic states may be observed near cancerous tissue regions.^47^

**Figure 6:**
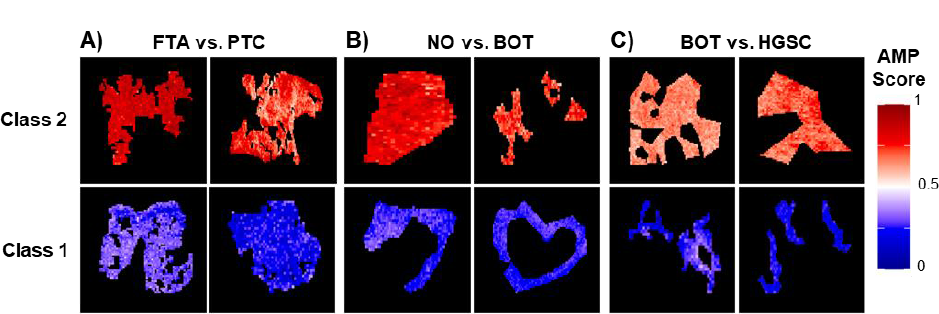
Example AMP score heatmaps for select A) FTA vs. PTC, B) NO vs. BOT, and C) BOT vs. HGSC homogenous tissues. For each tissue section, individual pixels are colored by their respective AMP score with lower scores shown in blue, midrange scores in white, and higher scores in red.

**Figure 7:**
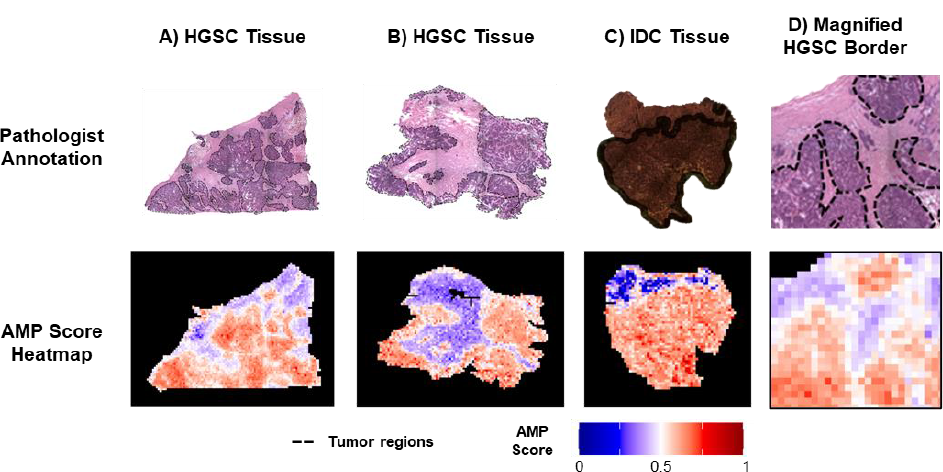
Comparison of AMP score heatmaps for heterogeneous tissues to annotated H&E slides for example HGSC tissue shown in **A)** and **B)**, IDC tissue in **C)**, and a magnified border region from **A)** in **D).** On the H&E slides, tissue outlined in black corresponds to tumorous areas with healthy tissue in the other regions.

## CONCLUSIONS

In this manuscript, we developed an AMP score pipeline to assess phenotypic differences in MSI data. We assessed these scores using data from three previous studies examining differences in normal, benign, and cancerous human tissue. Using the different studies, we first illustrated how AMP scores can improve tissue diagnostics by providing precise phenotypic predictions, as demonstrated by the high accuracy, sensitivity, and specificity observed across comparisons. In addition to providing accurate predictions, AMP score heatmaps highlight the utility of their continuous rather than categorical outcome. For example, at phenotype borders, intermediate AMP scores were observed, suggesting that these regions may have a combination of characteristics similar to both phenotypes. This finding provides insight into the molecular changes that take place as malignancies spread, as well as further informs decision making in surgical settings as physicians can choose to be more or less aggressive around these borders based on the nature of the disease.^48^ Beyond diagnostic utility, the AMP scores were also useful for visualizing tissue sections by weighting and summarizing significant features, thus circumventing the need to visualize each feature’s *m/z* map individually. Finally, since AMP scores are calculated before being combined with spatial information, the AMP scoring framework is also applicable to different biofluids such as urine and blood for patient classification. Thus, future work will include evaluating these different scenarios to define other AMP score use cases.^47^

## ASSOCIATED CONTENT

### Supporting Information

Overlap of selected features (**Figure S1**); ROC curves for phenotype predictions (**Figure S2**) (doc)

## AUTHOR INFORMATION

### Co-corresponding Author

The authors declare no competing interests.

## ACKNOWLEDGEMENTS

This work was funded by grants from the National Institute of Environmental Health Sciences (P42 ES027704 and P42 ES031009), the National Institute of General Medical Sciences (R01 GM141277), the National Institute of Health National Cancer Institute (R33 CA229068), the Welch Foundation (Q-1895-20220331), and a cooperative agreement with the Environmental Protection Agency (STAR RD 84003201). The views expressed in this manuscript do not reflect those of the funding agencies. We also would like to thank Marta Sans, Kyana Y. Garza, and Rachel J. DeHoog for their hard work collecting the data analyzed in this manuscript.

## REFERENCES

(1) Vaysse, P.-M.; Heeren, R. M. A.; Porta, T.; Balluff, B. Mass spectrometry imaging for clinical research – latest developments, applications, and current limitations. The Analyst 2017, 142 (15), 2690–2712. DOI: 10.1039/c7an00565b.

(2) Berghmans, E.; Boonen, K.; Maes, E.; Mertens, I.; Pauwels, P.; Baggerman, G. Implementation of MALDI Mass Spectrometry Imaging in Cancer Proteomics Research: Applications and Challenges. J Pers Med 2020, 10 (2). DOI: 10.3390/jpm10020054 From NLM PubMed-not-MEDLINE.

(3) Veta, M.; Pluim, J. P.; van Diest, P. J.; Viergever, M. A. Breast cancer histopathology image analysis: a review. IEEE Trans Biomed Eng 2014, 61 (5), 1400–1411. DOI: 10.1109/TBME.2014.2303852 From NLM Medline.

(4) Igbokwe, A.; Lopez-Terrada, D. H. Molecular testing of solid tumors. Arch Pathol Lab Med 2011, 135 (1), 67-82. DOI: 10.5858/2010-0413-RAR.1 From NLM Medline.

(5) Tian, H.; Sparvero, L. J.; Amoscato, A. A.; Bloom, A.; Bayir, H.; Kagan, V. E.; Winograd, N. Gas Cluster Ion Beam Time-of-Flight Secondary Ion Mass Spectrometry High-Resolution Imaging of Cardiolipin Speciation in the Brain: Identification of Molecular Losses after Traumatic Injury. Anal Chem 2017, 89 (8), 4611–4619. DOI: 10.1021/acs.analchem.7b00164 From NLM Medline.

(6) Ifa, D. R.; Eberlin, L. S. Ambient Ionization Mass Spectrometry for Cancer Diagnosis and Surgical Margin Evaluation. Clin Chem 2016, 62 (1), 111–123. DOI: 10.1373/clinchem.2014.237172 From NLM Medline.

(7) Ucal, Y.; Durer, Z. A.; Atak, H.; Kadioglu, E.; Sahin, B.; Coskun, A.; Baykal, A. T.; Ozpinar, A. Clinical applications of MALDI imaging technologies in cancer and neurodegenerative diseases. Biochim Biophys Acta Proteins Proteom 2017, 1865 (7), 795–816. DOI: 10.1016/j.bbapap.2017.01.005 From NLM Medline.

(8) Spraggins, J. M.; Rizzo, D. G.; Moore, J. L.; Noto, M. J.; Skaar, E. P.; Caprioli, R. M. Next-generation technologies for spatial proteomics: Integrating ultra-high speed MALDI-TOF and high mass resolution MALDI FTICR imaging mass spectrometry for protein analysis. Proteomics 2016, 16 (11-12), 1678–1689. DOI: 10.1002/pmic.201600003 From NLM Medline.

(9) Pagni, F.; De Sio, G.; Garancini, M.; Scardilli, M.; Chinello, C.; Smith, A. J.; Bono, F.; Leni, D.; Magni, F. Proteomics in thyroid cytopathology: Relevance of MALDI-imaging in distinguishing malignant from benign lesions. Proteomics 2016, 16 (11-12), 1775–1784. DOI: 10.1002/pmic.201500448 From NLM Medline.

(10) Wu, C.; Dill, A. L.; Eberlin, L. S.; Cooks, R. G.; Ifa, D. R. Mass spectrometry imaging under ambient conditions. Mass Spectrometry Reviews 2013, 32 (3), 218–243. DOI: 10.1002/mas.21360.

(11) Deininger, S.-O.; Ebert, M. P.; Fütterer, A.; Gerhard, M.; Röcken, C. MALDI Imaging Combined with Hierarchical Clustering as a New Tool for the Interpretation of Complex Human Cancers. Journal of Proteome Research 2008, 7 (12), 5230–5236. DOI: 10.1021/pr8005777.

(12) Abdelmoula, W. M.; Balluff, B.; Englert, S.; Dijkstra, J.; Reinders, M. J. T.; Walch, A.; McDonnell, L. A.; Lelieveldt, B. P. F. Data-driven identification of prognostic tumor subpopulations using spatially mapped t-SNE of mass spectrometry imaging data. Proceedings of the National Academy of Sciences 2016, 113 (43), 12244–12249. DOI: 10.1073/pnas.1510227113.

(13) Trindade, G. F.; Abel, M. L.; Lowe, C.; Tshulu, R.; Watts, J. F. A Time-of-Flight Secondary Ion Mass Spectrometry/Multivariate Analysis (ToF-SIMS/MVA) Approach To Identify Phase Segregation in Blends of Incompatible but Extremely Similar Resins. Anal Chem 2018, 90 (6), 3936–3941. DOI: 10.1021/acs.analchem.7b04877 From NLM PubMed-not-MEDLINE.

(14) Trede, D.; Schiffler, S.; Becker, M.; Wirtz, S.; Steinhorst, K.; Strehlow, J.; Aichler, M.; Kobarg, J. H.; Oetjen, J.; Dyatlov, A.;, et al. Exploring three-dimensional matrix-assisted laser desorption/ionization imaging mass spectrometry data: three-dimensional spatial segmentation of mouse kidney. Anal Chem 2012, 84 (14), 6079–6087. DOI: 10.1021/ac300673y From NLM Medline.

(15) Inglese, P.; McKenzie, J. S.; Mroz, A.; Kinross, J.; Veselkov, K.; Holmes, E.; Takats, Z.; Nicholson, J. K.; Glen, R. C. Deep learning and 3D-DESI imaging reveal the hidden metabolic heterogeneity of cancer. Chemical Science 2017, 8 (5), 3500–3511. DOI: 10.1039/c6sc03738k.

(16) Sarkari, S.; Kaddi, C. D.; Bennett, R. V.; Fernandez, F. M.; Wang, M. D. Comparison of clustering pipelines for the analysis of mass spectrometry imaging data. 2014, IEEE. DOI: 10.1109/embc.2014.6944691.

(17) Prasad, M.; Postma, G.; Franceschi, P.; Buydens, L. M. C.; Jansen, J. J. Evaluation and comparison of unsupervised methods for the extraction of spatial patterns from mass spectrometry imaging data (MSI). Scientific Reports 2022, 12 (1). DOI: 10.1038/s41598-022-19365-4.

(18) Bemis, K. D.; Harry, A.; Eberlin, L. S.; Ferreira, C.; van de Ven, S. M.; Mallick, P.; Stolowitz, M.; Vitek, O. Cardinal: an R package for statistical analysis of mass spectrometry-based imaging experiments. Bioinformatics 2015, 31 (14), 2418–2420. DOI: 10.1093/bioinformatics/btv146 From NLM Medline.

(19) Vijayalakshmi, K.; Shankar, V.; Bain, R. M.; Nolley, R.; Sonn, G. A.; Kao, C. S.; Zhao, H.; Tibshirani, R.; Zare, R. N.; Brooks, J. D. Identification of diagnostic metabolic signatures in clear cell renal cell carcinoma using mass spectrometry imaging. International Journal of Cancer 2020, 147 (1), 256–265. DOI: 10.1002/ijc.32843.

(20) Lee, J. W.; Figeys, D.; Vasilescu, J. Biomarker assay translation from discovery to clinical studies in cancer drug development: quantification of emerging protein biomarkers. Adv Cancer Res 2007, 96, 269–298. DOI: 10.1016/S0065-230X(06)96010-2 From NLM Medline.

(21) Pell, R.; Oien, K.; Robinson, M.; Pitman, H.; Rajpoot, N.; Rittscher, J.; Snead, D.; Verrill, C.; Driskell, O. J.; Hall, A.;, et al. The use of digital pathology and image analysis in clinical trials. The Journal of Pathology: Clinical Research 2019, 5 (2), 81–90. DOI: 10.1002/cjp2.127.

(22) Eberlin, L. S.; Norton, I.; Dill, A. L.; Golby, A. J.; Ligon, K. L.; Santagata, S.; Cooks, R. G.; Agar, N. Y. R. Classifying Human Brain Tumors by Lipid Imaging with Mass Spectrometry. Cancer Research 2012, 72 (3), 645–654. DOI: 10.1158/0008-5472.can-11-2465.

(23) Porcari, A. M.; Zhang, J.; Garza, K. Y.; Rodrigues-Peres, R. M.; Lin, J. Q.; Young, J. H.; Tibshirani, R.; Nagi, C.; Paiva, G. R.; Carter, S. A.;, et al. Multicenter Study Using Desorption-Electrospray-Ionization-Mass-Spectrometry Imaging for Breast-Cancer Diagnosis. Anal Chem 2018, 90 (19), 11324–11332. DOI: 10.1021/acs.analchem.8b01961 From NLM Medline.

(24) Guenther, S.; Muirhead, L. J.; Speller, A. V.; Golf, O.; Strittmatter, N.; Ramakrishnan, R.; Goldin, R. D.; Jones, E.; Veselkov, K.; Nicholson, J.;, et al. Spatially resolved metabolic phenotyping of breast cancer by desorption electrospray ionization mass spectrometry. Cancer Res 2015, 75 (9), 1828–1837. DOI: 10.1158/0008-5472.CAN-14-2258 From NLM Medline.

(25) DeHoog, R. J.; Zhang, J.; Alore, E.; Lin, J. Q.; Yu, W.; Woody, S.; Almendariz, C.; Lin, M.; Engelsman, A. F.; Sidhu, S. B.;, et al. Preoperative metabolic classification of thyroid nodules using mass spectrometry imaging of fine-needle aspiration biopsies. Proc Natl Acad Sci U S A 2019, 116 (43), 21401–21408. DOI: 10.1073/pnas.1911333116 From NLM Medline.

(26) Eberlin, L. S.; Tibshirani, R. J.; Zhang, J.; Longacre, T. A.; Berry, G. J.; Bingham, D. B.; Norton, J. A.; Zare, R. N.; Poultsides, G. A. Molecular assessment of surgical-resection margins of gastric cancer by mass-spectrometric imaging. Proc Natl Acad Sci U S A 2014, 111 (7), 2436–2441. DOI: 10.1073/pnas.1400274111 From NLM Medline.

(27) Sans, M.; Gharpure, K.; Tibshirani, R.; Zhang, J.; Liang, L.; Liu, J.; Young, J. H.; Dood, R. L.; Sood, A. K.; Eberlin, L. S. Metabolic Markers and Statistical Prediction of Serous Ovarian Cancer Aggressiveness by Ambient Ionization Mass Spectrometry Imaging. Cancer Res 2017, 77 (11), 2903–2913. DOI: 10.1158/0008-5472.CAN-16-3044 From NLM Medline.

(28) Eberlin, L. S.; Dill, A. L.; Costa, A. B.; Ifa, D. R.; Cheng, L.; Masterson, T.; Koch, M.; Ratliff, T. L.; Cooks, R. G. Cholesterol sulfate imaging in human prostate cancer tissue by desorption electrospray ionization mass spectrometry. Anal Chem 2010, 82 (9), 3430–3434. DOI: 10.1021/ac9029482 From NLM Medline.

(29) Dill, A. L.; Eberlin, L. S.; Zheng, C.; Costa, A. B.; Ifa, D. R.; Cheng, L.; Masterson, T. A.; Koch, M. O.; Vitek, O.; Cooks, R. G. Multivariate statistical differentiation of renal cell carcinomas based on lipidomic analysis by ambient ionization imaging mass spectrometry. Anal Bioanal Chem 2010, 398 (7-8), 2969–2978. DOI: 10.1007/s00216-010-4259-6 From NLM Medline.

(30) Paine, M. R.; Kim, J.; Bennett, R. V.; Parry, R. M.; Gaul, D. A.; Wang, M. D.; Matzuk, M. M.; Fernandez, F. M. Whole Reproductive System Non-Negative Matrix Factorization Mass Spectrometry Imaging of an Early-Stage Ovarian Cancer Mouse Model. PLoS One 2016, 11 (5), e0154837. DOI: 10.1371/journal.pone.0154837 From NLM Medline.

(31) Eberlin, L. S.; Margulis, K.; Planell-Mendez, I.; Zare, R. N.; Tibshirani, R.; Longacre, T. A.; Jalali, M.; Norton, J. A.; Poultsides, G. A. Pancreatic Cancer Surgical Resection Margins: Molecular Assessment by Mass Spectrometry Imaging. PLOS Medicine 2016, 13 (8), e1002108. DOI: 10.1371/journal.pmed.1002108.

(32) Gu, J.; Taylor, C. R. Practicing Pathology in the Era of Big Data and Personalized Medicine. Applied Immunohistochemistry &amp; Molecular Morphology 2014, 22 (1), 1–9. DOI: 10.1097/pai.0000000000000022.

(33) Tibshirani, R. Regression Shrinkage and Selection via the Lasso. Journal of the Royal Statistical Society 1996, 58 (1), 267–288.

(34) Friedman, J.; Hastie, T.; Tibshirani, R. Regularization Paths for Generalized Linear Models via Coordinate Descent. J Stat Softw 2010, 33 (1), 1–22. From NLM PubMed-not-MEDLINE.

(35) Breiman, L. Random forests. Machine Learning 2001, 45 (1), 5–32. DOI: 10.1023/a:1010933404324.

(36) Liaw A, W. M. Classification and Regression by randomForest. In 2002.

(37) Boser, B. E.; Guyon, I. M.; Vapnik, V. N. A training algorithm for optimal margin classifiers. 1992, ACM. DOI: 10.1145/130385.130401.

(38) Pedregosa, F. a. V., Ga{\"e}l and Gramfort, Alexandre and Michel, Vincent and Thirion, Bertrand and Grisel, Olivier and Blondel, Mathieu and Prettenhofer, Peter and Weiss, Ron and Dubourg, Vincent and others. Scikit-learn: Machine learning in Python. Journal of machine learning research 2011, 12, 2825–2830.

(39) Youden, W. J. Index for rating diagnostic tests. Cancer 1950, 3 (1), 32–35. DOI: 10.1002/1097-0142(1950)3:1&lt;32::aid-cncr2820030106>3.0.co;2-3 From NLM Medline.

(40) Hirschfeld, C. T. a. G. cutpointr : Improved Estimation and Validation of Optimal Cutpoints in R. Journal of Statistical Software 2021, 98. DOI: 10.18637/jss.v098.i11.

(41) Robin, X.; Turck, N.; Hainard, A.; Tiberti, N.; Lisacek, F.; Sanchez, J.-C.; Müller, M. pROC: an open-source package for R and S+ to analyze and compare ROC curves. BMC Bioinformatics 2011, 12 (1), 77. DOI: 10.1186/1471-2105-12-77.

(42) Wickham, H. ggplot2: Elegant Graphics for Data Analysis. Springer-Verlag New York 2016.

(43) Pes, B. Ensemble feature selection for high-dimensional data: a stability analysis across multiple domains. Neural Computing and Applications 2020, 32 (10), 5951–5973. DOI: 10.1007/s00521-019-04082-3.

(44) Bagley, S. C.; White, H.; Golomb, B. A. Logistic regression in the medical literature: standards for use and reporting, with particular attention to one medical domain. J Clin Epidemiol 2001, 54 (10), 979–985. DOI: 10.1016/s0895-4356(01)00372-9 From NLM Medline.

(45) Ruopp, M. D.; Perkins, N. J.; Whitcomb, B. W.; Schisterman, E. F. Youden Index and optimal cut-point estimated from observations affected by a lower limit of detection. Biom J 2008, 50 (3), 419–430. DOI: 10.1002/bimj.200710415 From NLM Medline.

(46) Cao, X.; Ding, L.; Mersha, T. B. Development and validation of an RNA-seq-based transcriptomic risk score for asthma. Scientific Reports 2022, 12 (1). DOI: 10.1038/s41598-022-12199-0.

(47) Woolman, M.; Katz, L.; Gopinath, G.; Kiyota, T.; Kuzan-Fischer, C. M.; Ferry, I.; Zaidi, M.; Peters, K.; Aman, A.; McKee, T.;, et al. Mass Spectrometry Imaging Reveals a Gradient of Cancer-like Metabolic States in the Vicinity of Cancer Not Seen in Morphometric Margins from Microscopy. Anal Chem 2021, 93 (10), 4408–4416. DOI: 10.1021/acs.analchem.0c04129 From NLM Medline.

(48) Batlle, E.; Wilkinson, D. G. Molecular Mechanisms of Cell Segregation and Boundary Formation in Development and Tumorigenesis. Cold Spring Harbor Perspectives in Biology 2012, 4 (1), a008227–a008227. DOI: 10.1101/cshperspect.a008227.

