## Supplemental Figures for "Utilizing Aggregated Molecular Phenotype (AMP) Scores to Visualize Simultaneous Molecular Changes in Mass Spectrometry Imaging Data"

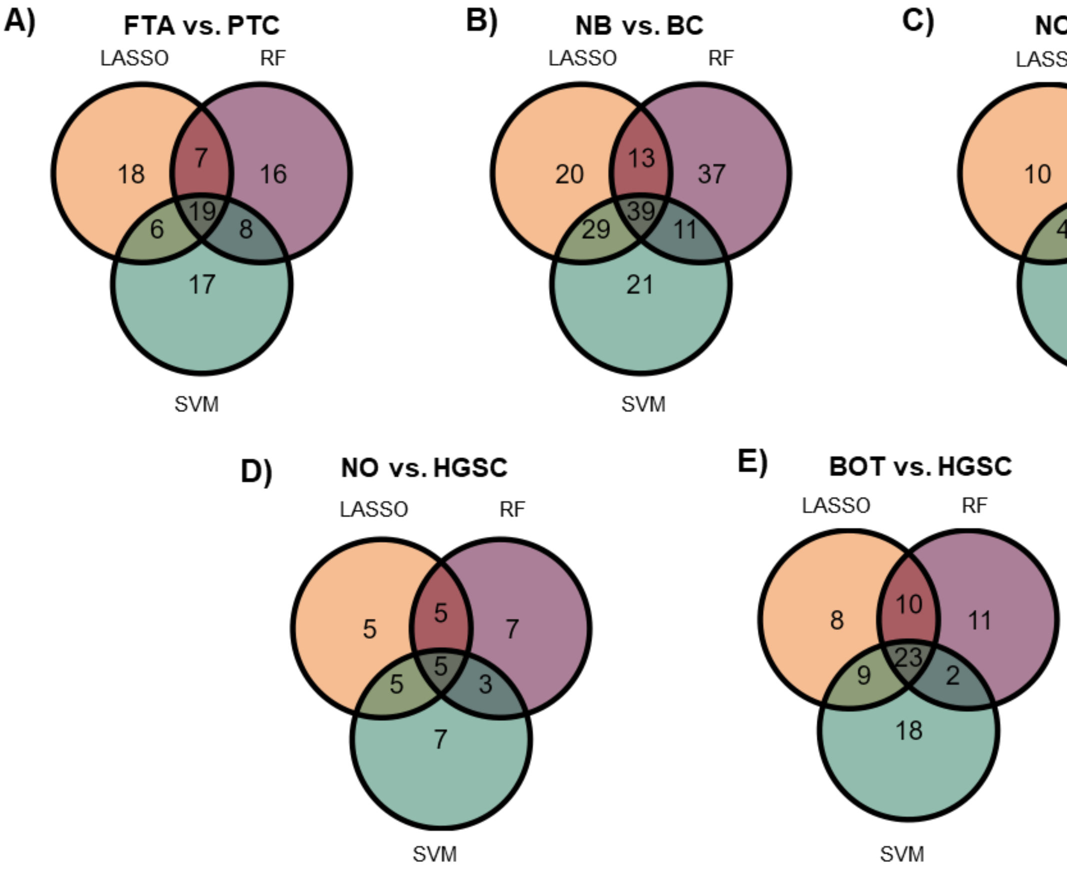


**Figure S1:** Venn diagrams showing the overlap of features selected by Lasso, random forest (RF) and support vector machine (SVM) for A) FTA vs. PTC samples, B) NB vs. BC samples, C) NO vs. BOT samples, D) NO vs. HGSC, and E) BOT vs. HGSC samples.

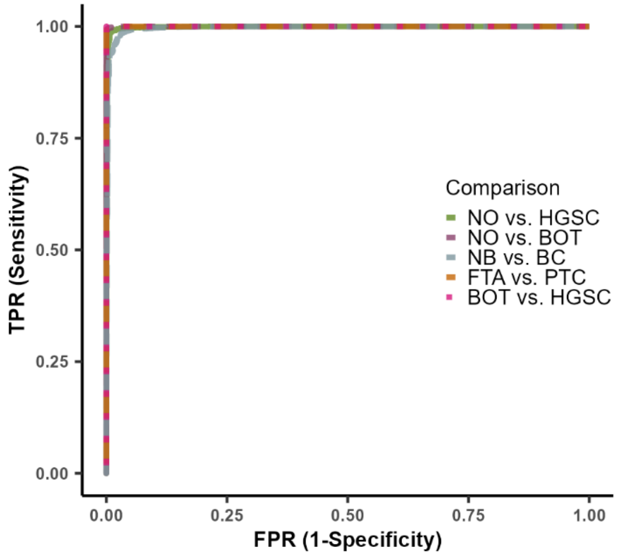


**Figure S2:** Receiver operator characteristic (ROC) curves for phenotype prediction performance of AMP scores for each pairwise comparison.
